# Evidence of an acetone carboxylation pathway in photoheterotrophic bacteria from the Arctic Ocean

**DOI:** 10.64898/2026.08.16.745133

**Authors:** Susan McLatchie, Sara Palestini, Asher Woodhead, Tony Gutierrez, David A. Walsh

## Abstract

Carboxylases are among the most important enzymes in nature as they catalyze the fixation of inorganic carbon (CO_2_), a central step in the global carbon cycle. In addition to their well-known function in autotrophic CO_2_ fixation, many carboxylases play a role in the heterotrophic assimilation of organic compounds. In this study, we provide genomic evidence for an assimilatory carboxylation pathway involved in acetone degradation in photoheterotrophic bacteria from metagenomes collected along a latitudinal transect of the Arctic Ocean. This curious metabolism was linked to a single population of Gammaproteobacteria (*Porticoccus arcticus*). *P. arcticus* has a streamlined genome compared to *Porticoccus* relatives but has maintained a complete acetone carboxylation pathway while acquiring multiple proteorhodopsin genes by lateral gene transfer. Arctic Ocean metatranscriptomes revealed the acetone carboxylase and rhodopsins genes were among the most highly expressed *P. arcticus* genes in oligotrophic Arctic surface waters. *P. arcticus* sequences were consistently detected, and often abundant (up to 9%), in a multiyear Arctic Ocean 16S rRNA time-series, supporting its ecological significance in Arctic marine systems. Overall, this work reports a metabolic module (acetone carboxylation) in the ocean that may allow photoheterotrophic bacteria to enhance their biosynthetic capacity via CO_2_ assimilation.

## Introduction

Carboxylation reactions are among the most important in nature because they lead to the fixation of inorganic carbon (CO_2_), which is central to the global carbon cycle^1^. Carboxylases play a role in diverse microbial metabolisms such as autotrophy, anaplerosis, and heterotrophic carbon assimilation^1^. Photosynthesis is mediated through autotrophic carboxylases and is the dominant carbon fixation process in the sunlit ocean, while anaplerotic carboxylases are involved in the replenishment of citric acid cycle intermediates. Assimilatory carboxylases function in heterotrophic pathways that convert organic growth substrates into central metabolic intermediates^1^. Compared to autotrophic and anaplerotic carboxylases, the diversity and ecological role of assimilatory carboxylases are generally much less understood. Nevertheless, relatively high rates of dark CO_2_ assimilation linked to heterotrophs have been reported in the ocean, particularly under oligotrophic conditions^2,3^.

One assimilatory carboxylase that may be significant in marine bacteria is associated with acetone degradation. Acetone degradation via carboxylation was first discovered and characterized in terrestrial bacteria^4,5^. The first step is carried out by an ATP-dependent acetone carboxylase (AcxABC) that converts acetone to acetoacetate^6^. Following carboxylation, acetoacetate is converted via two enzymatic reactions to acetyl-CoA, which feeds into central metabolism. Acetone is a common volatile organic compound in the ocean^7^ with sources that include phytoplankton production^8^, photochemical degradation of organic matter^9^, and atmospheric deposition^10^. Biological consumption of acetone in ocean surface waters has been reported^11^. Oligotrophic marine bacteria (i.e. SAR11) can consume acetone although the acetone degradation proceeds through an acetone monooxygenase-mediated pathway^12^. Similar monooxygenases are common in genomes of diverse marine bacteria and metatranscriptomes, supporting the notion that acetone is an important carbon source in the ocean^12^. However, acetone degradation via carboxylation has yet to be described in marine bacteria. Acetone carboxylation may represent a previously underappreciated carbon assimilation pathway advantageous to marine bacteria living under organic carbon-limited conditions.

The aim of this study was to identify and characterize marine bacteria with the metabolic potential for carbon assimilation via acetone carboxylation. We focused our investigation on the Arctic Ocean (Canada Basin) since previous work has shown that Arctic Ocean surface water has a relatively high concentration of acetone (> 40 nM acetone in surface waters)^13^, acts as a biological sink for acetone^14^, is a location of significant heterotrophic CO_2_ assimilation^15^, and the surface waters are oligotrophic^16^. Given this biogeochemical setting, we hypothesized that heterotrophic Arctic Ocean bacteria may benefit from acetone-mediated CO_2_ assimilation.

In this work, we provide genomic evidence for an assimilatory carboxylation pathway involved in acetone degradation in photoheterotrophic bacteria from Arctic Ocean metagenomes. We identify this metabolism in a single population of Gammaproteobacteria (*Porticoccus arcticus*). We show that *P. arcticus* has a streamlined genome compared to *Porticoccus* relatives but has maintained a complete acetone carboxylation pathway while acquiring multiple proteorhodopsin genes by lateral gene transfer. Arctic Ocean metatranscriptomic surveys reveal the acetone carboxylase and rhodopsins genes are among the most highly expressed *P. arcticus* genes in oligotrophic Arctic surface waters. Overall, this work reports on an acetone carboxylation pathway in the ocean that may allow photoheterotrophic bacteria to enhance their biosynthetic capacity via CO_2_ assimilation.

## Results

### Acetone-dependent growth in a marine bacterium

We first sought evidence for the presence of the acetone carboxylation pathway in cultured marine bacteria. A complete operon for acetone carboxylase (*acxABC*) was identified in the genome of *Porticoccus hydrocarbonoclasticus* strain MCTG13d, a bacterium previously isolated from a dinoflagellate laboratory culture^17^. In addition to *acxABC*, the *P. hydrocarbonoclasticus* genome encoded the succinyl-CoA:acetoacetate transferase (*scoAB*) and acetyl-CoA acetyltransferase (*atoD*) enzymes necessary for transformation of acetoacetate to acetyl-CoA (**Figure 1a**). Acetone metabolism by *P. hydrocarbonoclasticus* was then tested. Acetone provided as a sole source of organic carbon supported *P. hydrocarbonoclasticus* growth (**Figure 1b**), demonstrating that marine bacteria carrying the *acxABC* genes can metabolize and use acetone for growth. Interestingly, growth rate and maximum cell density were inversely related to initial acetone concentration. We posit this is likely due to increasing toxicity to the cells with increasing acetone concentration.

**Figure 1.**
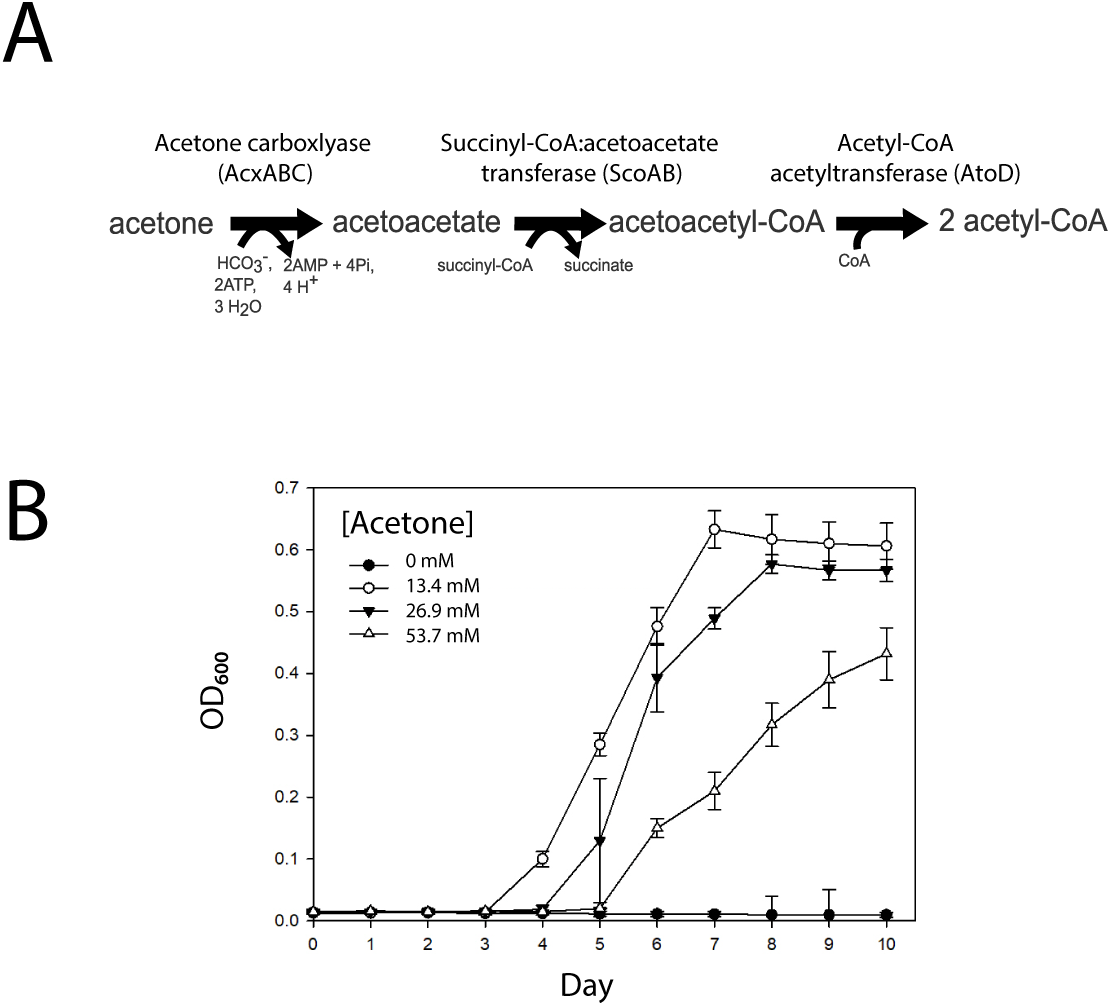
An acetone degradation pathway in marine bacteria. **(A)** The acetone degradation pathway identified in the genome of *P. hydrocarbonoclasticus* strain MCTG13d (**B**) Growth curves for P. hydrocarbonoclasticus strain MCTG13d grown on acetone concentrations (13.4 mM, 26.9 mM, 53.7 mM) as the sole source of carbon compared to non-amended (no acetone) control. Five replicates were used for each of the three acetone concentrations. Error bars represent standard deviation. Source data are provided as a source data file.

### Diversity and expression of acetone carboxylase in the Arctic Ocean

Using the acetone carboxylase alpha subunit *(acxB)* as a functional gene marker, we searched an Arctic Ocean (Canada Basin) metagenome-assembled genome (MAG) catalogue for evidence of acetone carboxylation (see Supplementary Material for further details on MAG catalogue generation, **Figure S1, Figure S2**). Fifteen putative *acxB* genes were identified in the MAG catalogue (encoded across 12 MAGs). Acetone carboxylase is a member of the larger hydantoinase/oxoprolinase protein family, which contains enzymes with diverse substrate specificities beyond acetone^18^. To increase confidence that *acxB* homologs from the Arctic Ocean MAGs encoded an acetone carboxylase enzyme, we reconstructed the phylogenetic relationship between AcxB identified in the MAG catalogue and genes encoding known acetone carboxylase as well as hydantoinase and acetophenone carboxylase (**Figure 2a**). Two homologs (Port-CB9S-26_00193 and Gamma-CB-1185_00243) were positioned within a well-supported clade that contained experimentally validated acetone carboxylase enzymes and the enzyme from *P. hydrocarbonoclasticus*. The remaining 13 homologs were positioned across three deeper branching clades. Although these clades were more closely related to known acetone carboxylases compared to hydantoinase and acetophenone carboxylase, we are less confident in their substrate specificity.

**Figure 2.**
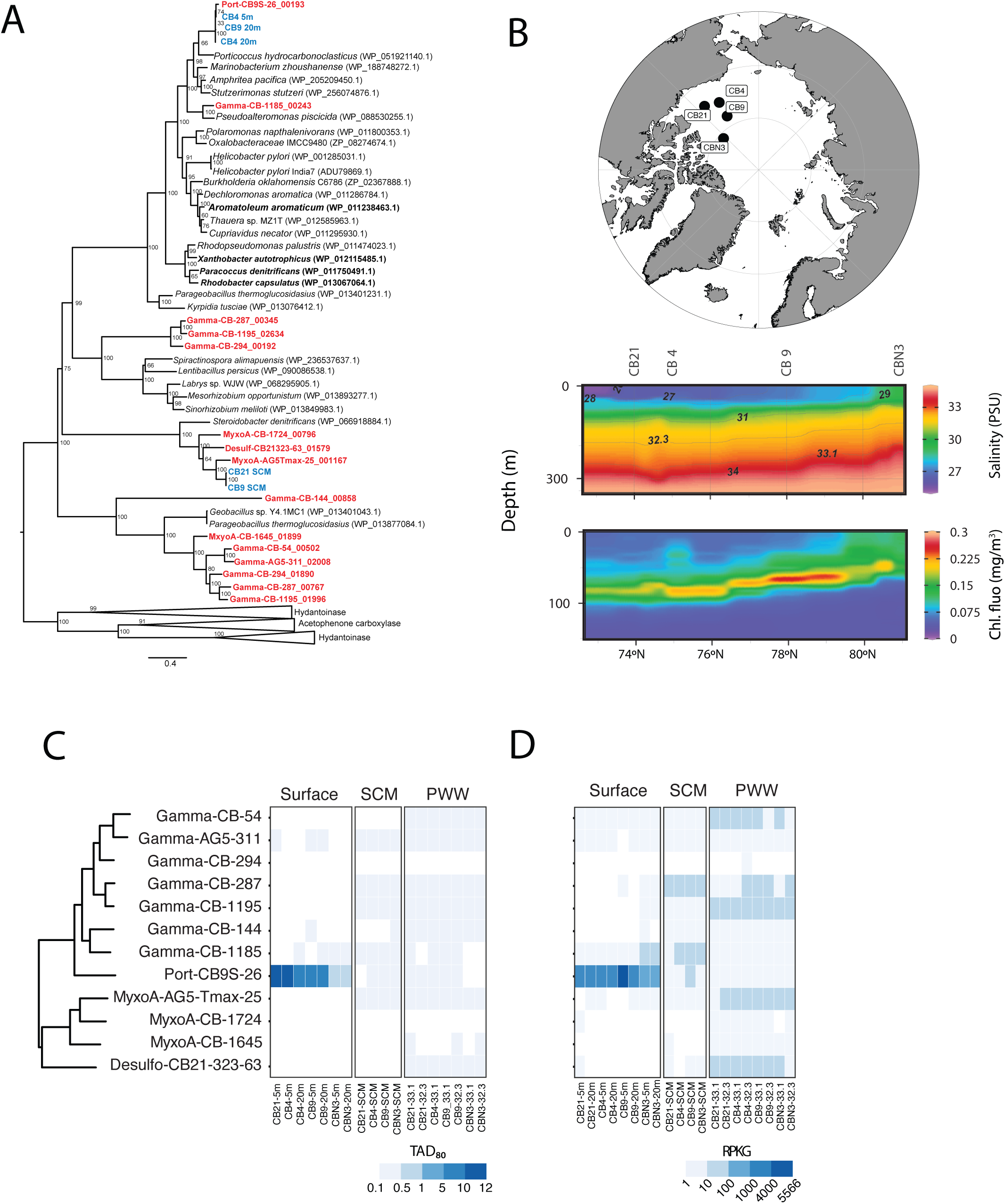
AcxB surveys reveal acetone degradation pathways in Arctic Ocean microbiomes. (**A**) Phylogenetic relationships between acetone carboxylase *acxB* gene from the Canada Basin MAG dataset (red) and the Canada Basin metagenome assemblies (blue). Experimentally validated acetone-consuming bacteria are highlighted in bold. A maximum likelihood tree was constructed from protein sequence data using the JTT substitution model, gamma distribution (4 categories), and the nearest neighbor heuristic search model with 100 bootstrap replications. The tree was rooted using the alpha subunit gene of distantly related members of the hydantoinase/oxoprolinase protein family. Taxa in bold are those in which acetone carboxylase activity has been experimentally validated. (**B**) Locations and environmental profiles for samples collected in September 2017 and used in metagenomic and metatranscriptomic fragment recruitment. The map was generated using ggOceanMaps in R^42^. (**C**) Abundance of MAGs encoding an *acxB* gene based on fragment recruitment of metagenomic reads. MAG names reflect taxonomy from GTDB phylogenomic classification. (**D**) Representation of MAGs in metatranscriptomic data based on fragment recruitment of metatranscriptomic reads. Source data are provided as a source data file.

To further explore AcxB diversity in the Arctic Ocean, we expanded the search for AcxB homologs beyond MAGs to a collection of metagenome assemblies from the vertically stratified Arctic Ocean. Metagenome assemblies represented the surface waters (Surface; 5 m and 20 m), the subsurface chlorophyll maximum (SCM; 55-95 m), and deeper Pacific-origin winter waters (PWW; 90-250 m). We retrieved five additional acxB homologs from metagenome assemblies that were not previously identified in the MAGs. Three AcxB homologs identified in surface metagenomes shared 98% amino acid identity with the Port-CB9S-26_00193 homolog, while two AcxB homologs identified in SCM metagenomes were positioned in one of the more distantly related clades (**Figure 2a**). These results demonstrate a limited diversity of *acxB* genes in the Arctic Ocean and that the MAGs represent the dominant *acxB*-containing assemblage.

We next investigated the abundance and distribution of the AcxB-containing MAGs across the Arctic Ocean (**Figure 2b**). Metagenomic fragment recruitment revealed that a single MAG taxonomically assigned to the *Porticoccaceae* (Port-CB9S-26) dominated the AcxB-containing assemblages in ice-free surface waters (up to 12 TAD_80_) (**Figure 2c**). Port-CB9S-26 was less abundant at the ice-covered northern station CBN3 (0.8-0.9 TAD_80_) and found at very low abundance in the SCM and PWW. The other 11 MAGs were relatively rare (< 0.3 TAD_80_) and were more common in deeper water layers.

In addition to the metagenomic survey, we performed an overlapping metatranscriptomic survey of the Arctic Ocean^19^, allowing the assessment of MAG gene expression abundance patterns. Metatranscriptomic fragment recruitment to MAGs revealed Port-CB9S-26 dominated the AcxB-containing assemblages in surface water metatranscriptomes (387-5566 RPKG) (**Figure 2d**). The other 11 MAGs were much less represented in metatranscriptomes and were more common in deeper water layers. These results demonstrate that the abundant and transcriptionally active Port-CB9S-26 MAG represents the major population of putative acetone-carboxylating bacteria in the Arctic Ocean surface waters.

### Phylogenetic identity and comparative genomics

The Port-CB9S-26 MAG encoded a single complete rRNA operon and 16S rRNA phylogenetic analysis confirmed taxonomic assignment within *Porticoccaceae* (**Figure 3a**). Herein, we refer to the bacterial population represented by the Port-CB9S-26 MAG as *Porticoccus arcticus. P. arcticus* formed a clade with environmental 16S rRNA sequences from coastal waters, suggesting a broader oceanic distribution. An additional member of this clade was *Porticoccus* sp. Uisw_050_02, which was previously identified in a mixed enrichment culture from the Arctic Ocean^20^. The *P. arcticus* genome (1.29 Mbp, 41.6 %GC) and *Porticoccus* sp. Uisw_050_02 genome (1.44 Mbp, 41.5 %GC) shared a similar small size and lower GC content compared to the publicly available *Porticoccaceae* genomes of *P. hydrocarbonoclasticus* (2.47 Mbp, 53.1% GC) (**Figure 3b**) and the SAR92 strain HTCC2207 (2.63 Mbp, 49.1% GC). Comparative genomics revealed that 98% (1179 of 1207) of *P. arcticus* protein-coding genes are shared with *Porticoccus* sp. Uisw_050_02. The *Porticoccus* sp. Uisw_050_02 genome is complete (a single closed circular chromosome) and carries an additional 174 protein-coding genes compared to *P. arcticus,* which may reflect the incomplete nature of the MAG or variation in gene content. Overall, the relatively small genome size and lower GC content suggests *P. arcticus* evolution has involved genome reduction.

**Figure 3.**
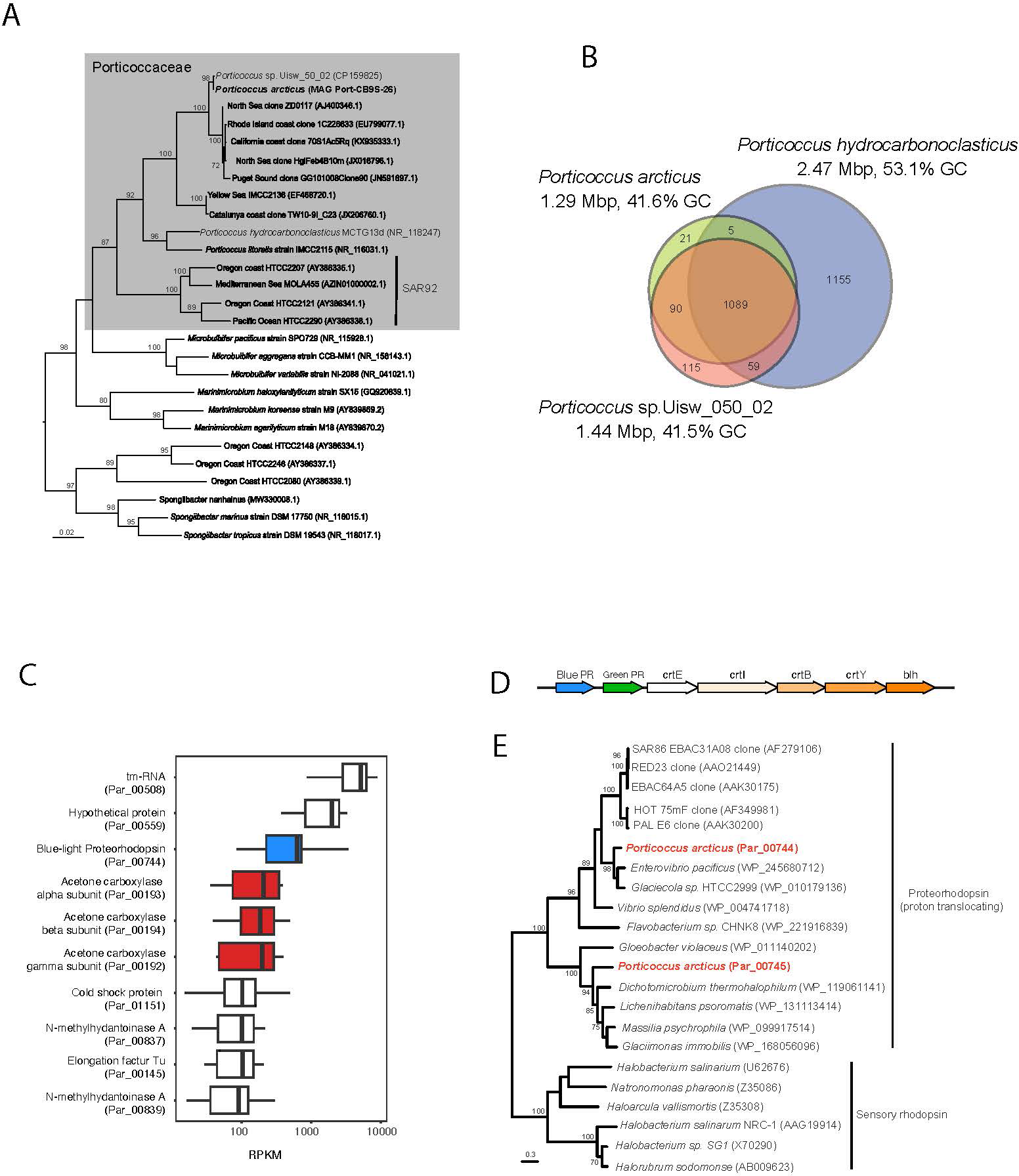
Phylogenetic identification and metabolic characteristics of *Porticoccus arcticus*. (**A**) Maximum likelihood phylogeny of marine Gammaproteobacteria based on 16S rRNA gene sequences. The tree was constructed using the GTR + gamma distribution (4 categories) model of nucleotide substitution in MEGA v.11 with 100 bootstraps. (**B**) Genomic characteristics and distribution of genes between *P. arcticus, Porticoccus* sp. Uisw_50_02, and *P. hydrocarbonoclasticus*. Values in the Venn diagram are the number of genes. (**C**) The ten most abundant *P. arcticus* transcripts identified in Canada Basin surface water metatranscriptomes. Boxes represent the interquartile range (IQR; 25th–75th percentiles), the horizontal line indicates the median, whiskers extend to 1.5 × IQR calculated from a total of 8 metatranscriptomes. Acetone carboxylase genes are highlighted in red and proteorhodopsin is highlighted in blue. (**D**) Region within the *P. arcticus* genomes encoding the multiple proteorhodopsins genes and carotenoid biosynthesis genes. **(E)** Phylogenetic analysis of rhodopsin genes and the proteorhodopsin biosynthesis operon within the *P. arcticus* genome. A maximum likelihood tree was constructed using the JTT substitution model, gamma distribution (4 categories), and the nearest neighbor heuristic search model with 100 bootstrap replications. Bootstrap values less than 70 were removed. *P. arcticus* homologs are highlighted in red. Source data are provided as a source data file.

### Photoheterotrophic metabolism in *P. arcticus*

Metabolic reconstruction of *P. arcticus* revealed a specialized carbon metabolism. In addition to *acxABC*, the succinyl-CoA:acetoacetate transferase (*scoAB*) and acetyl-CoA acetyltransferase enzyme (*atoD*) necessary for transformation of acetone to acetyl-CoA were present (**Figure S3, Supplementary Data 4**). Expression of the full pathway was detected in metatranscriptomes and the *acxABC* subunits were among the top six most abundant transcripts (13-497 RPKM values) (**Figure 3c, Supplementary Data 5**). The genome encoded a cytoplasmic carbonic anhydrase, providing further evidence for CO_2_ as a metabolic substrate. Very few transporters for organic carbon compounds (3 TRAP and 2 oligopeptide transporters) were identified in the genome and no annotated sugar transporters were identified. Like *P. hydrocarbonoclasticus,* both *P. arcticus* and *Porticoccus* sp. Uisw_050_02 genomes lack glucokinase suggesting an incomplete glycolytic pathway, and the oxidative pentose phosphate pathway was also incomplete (**Figure S3**). Overall, these observations suggest a significant role for acetone as a growth substrate for *P. arcticus* and, like *P. hydrocarbonoclasticus* ^17^, an inability to use sugars as growth substrates.

Given the restricted spectrum of carbon compounds available to *P. arcticus*, we looked for additional energy generation pathways. The presence of two adjacent proteorhodopsin genes in *P. arcticus* suggested a capacity for phototrophic energy metabolism (**Figure 3d, Supplementary Data 4**). Both proteorhodopsins were predicted proton pumps based on phylogenetic position within the proteorhodopsin clade and amino acid identities at positions 97 (aspartate), 101 (threonine), and 108 (glutamate)^21^. One homolog was most closely related to proteorhodopsins from distantly related *Gammaproteobacteria* while the other was related to proteorhodopsins from *Betaproteobacteria*, suggesting both were acquired by lateral gene transfer (**Figure 3e**).

The proteorhodopsins were predicted to have distinct light absorbing characteristics. One was predicted to be blue light absorbing (presence of glutamate at position 105), while the other was green light absorbing (presence of leucine at position 105)^21^. The blue light sensing proteorhodopsin was one of the most highly expressed *P. arcticus* genes in metatranscriptomes (**Figure 3d**), while the green light absorbing proteorhodopsin was much less expressed (∼64-fold lower expression) (**Table S5**). A complete carotenoid biosynthesis pathway was located immediately downstream of the proteorhodopsin genes (**Figure 3d**). Expression of the complete carotenoid biosynthesis pathway was observed in the metatranscriptomes (**Supplementary Data 5**). Overall, these observations strongly support a photoheterotrophic metabolism specialized to the use of acetone as a carbon source in *P. arcticus*.

### Biogeography of *P. arcticus* in the Arctic Ocean

To further elucidate the biogeography of *P. arcticus*, we analyzed a multiyear 16S rRNA time-series (2017-2023) generated along a latitudinal gradient (72°N to 83°N) during the late summer/early autumn. The gradient comprised a consistently ice-free ocean site (BL8), multiple open ocean sites characterized by dynamic interannual ice-conditions (CB21, CB4, and CB9) and northern sites that were consistently ice-covered in summer (Ice stations) (**Figure 4, Supplementary Data 3**). A single amplicon sequence variant (ASV) was identified that exhibited 100% identity to the 16S rRNA gene in the *P. arcticus* genome. *P. arcticus* was detected in all 5 m and 20 m samples, indicating a broad distribution in Arctic Ocean surface waters. In contrast, *P. arcticus* was rarely detected in SCM samples. *P. arcticus* was relatively abundant in surface water samples from CB21, CB4, and CB9, representing up to 9 % of 16S rRNA sequences at 5 m and 20 m depths. In contrast, *P. arcticus* abundance was lower at the Ice stations and the consistently ice-free site. Overall, these results provide evidence that *P. arcticus* exhibits a habitat preference for oligotrophic Arctic surface waters with intermediate levels of sea ice cover.

**Figure 4.**
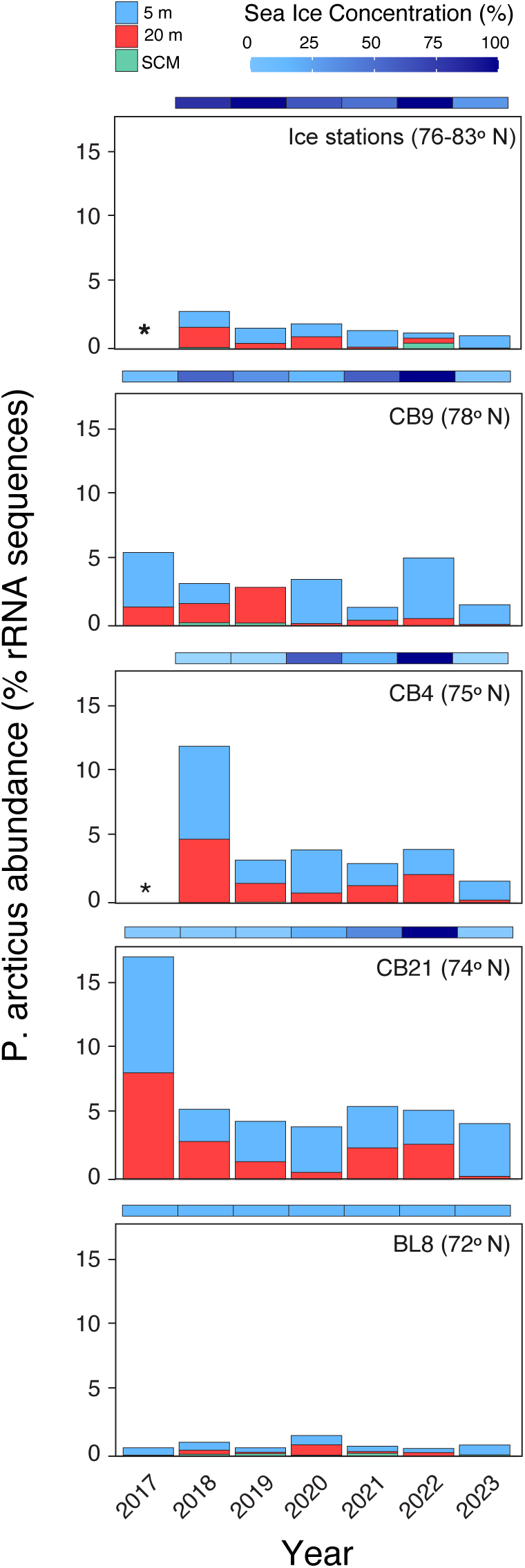
Distribution and abundance of *P. arcticus* in an Arctic Ocean 16S rRNA time-series. The sea ice concentration at the location and time of sampling is represented by the bar above each of the four plots. Asterisks represent sample locations where no data is available. Source data are provided as a source data file.

## Discussion

In this study, we identified and characterized a bacterial lineage (*P. arcticus*) that is widespread in the Arctic Ocean and exhibits remarkable adaptations to oligotrophic Arctic Ocean surface waters. *P. arcticus* has a streamlined genome compared to *Porticoccus* relatives but has maintained the metabolic capacity for acetone carboxylation while at the same time acquiring multiple proteorhodopsin genes via lateral gene transfer. In this way, *P. arcticus* has evolved a metabolic strategy that may be classified as a form of photoheterotrophy, where an otherwise heterotrophic metabolism is enhanced with a phototrophic energy metabolism and acetone-mediated CO_2_ assimilation. The acetone carboxylase and proteorhodopsin genes were among the most highly expressed in the genome, suggesting *P. arcticus* invests a large fraction of cellular resources into phototrophic energy conservation and acetone carboxylation. In doing so, this specialised metabolism may allow *P. arcticus* to take advantage of the sunlight and acetone that is available in ice-free summer surface waters of the Arctic Ocean, and to supplement its carbon requirement through CO_2_ assimilation via carboxylation.

*P. arcticus* was not only widespread in the Arctic Ocean but also reached relatively high abundances (nearly 10% of bacterioplankton) in surface waters. The surface samples in our study were from 5 and 20 m depths, which were located within the surface mixed layer. Previous work has shown a mean seawater acetone concentration of 9 nM, with elevated concentrations at the surface (>40 nM)^13^. Moreover, ultraviolet light (UV) is required for the photochemical production of acetone from chromophoric dissolved organic matter and UV light is rapidly absorbed within 2-7 m of the Arctic water column^22^. Hence, the association with acetone-enriched Arctic surface waters, and a general rarity at the SCM, fits with acetone playing a central role in *P. arcticus* metabolism. *P. arcticus* has multiple proteorhodopsins tuned to different wavelengths. A capacity to exploit light of different wavelength would be beneficial in the Arctic Ocean due to dynamic ice-cover and seasonal variability in the intensity and incidence angle of sunlight reaching the surface waters. Although our study focused on the late summer to early autumn period, we posit that *P. arcticus* may be more abundant earlier in the summer shortly after ice melt. The reduced sea ice cover may increase photochemical oxidation of organic matter in seawater^9,13^ and melting snowpack^23^, as well as increase air-sea exchange of acetone^24^, leading to an acetone pulse to surface water bacterial communities.

A notable observation from our study was that acetone carboxylation appears to be a rare metabolic trait in Arctic Ocean microbiomes. Other than *P. arcticus*, we only identified a few low abundance MAGs that contained the acetone carboxylase genes. Further analysis of the full metagenome assemblies did not lead to a significant increase in observed acetone carboxylase diversity. These observations suggest acetone degradation via carboxylation is a niche metabolism dominated by a single lineage of bacteria within the *Porticoccaceae*. One explanation is that there is competition for acetone with marine bacteria that possess the acetone monooxygenase-mediated pathway, which is widespread in the ocean^12^. Unlike acetone carboxylation, the monooxygenase-mediated pathway does not require an input of energy^25^ and is likely why it may be common in heterotrophic bacteria^12^ and possibly disseminated amongst marine bacteria via lateral gene transfer.

Another explanation for why the acetone carboxylation pathway has not been widely exchanged among marine bacteria is related to the concept of metabolic specialization promotion by biochemical conflict^26^. Cytoplasmic solvent capacity determines the maximum number of enzymes and other macromolecules that can be contained within a cell^27^. Cells that produce more enzymes for a specific pathway must produce fewer enzymes for other pathways and this competition for solvent capacity may promote specialization. Acetone carboxylase genes were among the most highly expressed in *P. arcticus*. We do not know if high transcript levels lead to high protein levels in *P. arcticus*, but previous work on acetone degrading isolates have shown a very high level of protein expression related to low specific activity of the enzyme^28^. If also true for *P. arcticus*, such a scenario may amount to a barrier to successful lateral gene transfer. Not only do the genes need to be transferred, but selection depends on enzyme expression levels. If more productive heterotrophic pathways are operating within the recipient populations, then maintenance of the acetone degradation pathway will be selected against through competition for cytoplasmic solvent capacity.

Overall, this study expands our understanding of bacterial diversity and adaptation to oligotrophic conditions in the Arctic Ocean, and more generally the known metabolic diversity of marine bacteria. Given the ongoing loss of sea ice, we speculate that *P. arcticus* may become a more prevalent and biogeochemically relevant member of a future Arctic Ocean microbiome. Future work characterising the global diversity of acetone carboxylases could illuminate whether this metabolism is in fact unique to the Arctic Ocean or more widespread in the global ocean.

## Methods

### Cultivation of P. hydrocarbonoclasticus

To assess the ability of *Porticoccus hydrocarbonoclasticus* strain MCTG13d to utilize acetone as a sole carbon and energy source, an inoculum of the organism was prepared by first growing colonies on ONR7a agar medium^29^ amended with 0.1% (w/v) pyruvate as the sole carbon source. Isolated colonies were inoculated into 10 mL ONR7a liquid medium amended with 0.1% (w/v) pyruvate. The cells were allowed to grow to the mid-to-late exponential phase, at which point the cells were pelleted (13,000 xg; 10 min) and washed three times with sterile ONR7a liquid medium. The washed cells were then resuspended in 0.5 mL of the same medium and used to inoculate a series of sterile, acid-washed (0.1 N HCl) Teflon-lined screw-cap glass tubes (100 × 13 mm), each containing 5 ml of synthetic seawater (ONR7a) medium amended with acetone as a sole organic carbon source to a final concentration of 13.4, 26.9 or 53.7 mM. Five replicates were used for each of the three acetone concentrations. All tubes were incubated with gentle rotary shaking (100 rpm) under natural daylight-night conditions (not direct sunlight) at 21°C, and growth was monitored spectrophotometrically at 600 nm over 2 weeks. Uninoculated medium and medium without added acetone (no carbon source) were used as controls.

### Identification and phylogenetic analysis of AxcB homologs

Acetone metabolism was identified in MAGs using the marker genes for acetone degradation acxABC (K10854, K10855, K10856). Reference sequences including those from experimentally validated acetone-consuming bacteria, full-length sequences the acetone carboxylase-harbouring MAGs, and sequences from the single metagenome assemblies (>500 bp in length) of the marker gene of acetone carboxylase (acxB) were included in a multiple sequence alignment using the MUSCLE algorithm in MEGA v.11^30^. The maximum likelihood tree was constructed using the JTT substitution model, gamma distribution (4 categories), and the nearest neighbor heuristic search model in MEGA v.11 with 100 bootstrap replicates^30^.

### Fragment recruitment

To determine the abundance of MAGs in the Canada Basin metagenomes, fragments were initially recruited to a concatenation of the MAGs at a minimum sequence identity of 98% using BBMap v.35. The mapping files were parsed to extract the mapping information for each individual MAG using a custom Unix script which implemented SAMtools^31^. Mapping files for each MAG were converted from SAM to sorted BAM format files using SAMtools^31^. Horizontal coverage was calculated following the method in^32^ in which the average sequencing depth is truncated to the central 80% of the mapped positions (TAD_80_) normalized by the number of genome equivalents.

To determine MAG gene expression abundance from, Canada Basin metatranscriptomes, we performed fragment recruitment of metatranscriptomic data previously described in^19^. Competitive fragment recruitment of the Canada Basin metatranscriptomes was carried out in the same manner as the metagenomic fragment recruitment. Metatranscriptomic RPKG values for the MAGs was calculated by dividing the total number of reads mapped to the MAG by the size of the MAG in Kbp and the size of the metatranscriptome in Gbp. For gene expression analysis in *P. arcticus*, metatranscriptomic RPKM values were calculated for each of the gene-coding sequences by dividing the total number of reads mapped to the gene by the size of the gene in Kbp and the total number of metatranscriptome reads in millions.

### 16S rRNA phylogeny

A multiple-sequence alignment of near full-length 16S rRNA genes from Gammaproteobacteria was generated using the MUSCLE algorithm as implemented in MEGA v.11^30^. A maximum likelihood tree was constructed using the GTR + gamma distribution (4 categories) model of nucleotide substitution in MEGA v.11 with 100 bootstraps^30^.

### Gene annotation and comparative genomics

Gene prediction and annotation was performed using Prokka v.1.12^33^, which implemented Prodigal v.2.6.3 for protein coding genes^34^, Barrnap for ribosomal RNA genes, and ARAGORN v.1.2^35^ for transfer and transfer-messenger RNA genes. Functional annotation of proteins was performed using KofamKOALA with default settings and a threshold score of 0.7 or higher and an e-value of 1 x 10^10^ or lower (Aramaki et al., 2020). To explore differences in gene content between *P. arcticus*, *Porticoccus* sp. Uisw_50_02, and *P. hydrocarbonoclasticus*, the distributions of orthologous genes were analyzed by Proteinortho^36^. The KEGG Mapper “Reconstruct” tool was used to identify central metabolic pathways harboured by *P. arcticus* and *P. hydrocarbonoclasticus*^37^.

### Identification and analysis of proteorhodopsins

To verify proteorhodopsin annotations, we performed a phylogenetic analysis to increase confidence in the biochemical function of the rhodopsin genes. Reference sequences for proteorhodopsin and sensory rhodopsin I and II were included in a multiple sequence alignment using the MUSCLE algorithm in MEGA v.11^30^. The maximum likelihood tree was constructed using the JTT substitution model, gamma distribution (4 categories), and the nearest neighbor heuristic search model in MEGA v.11 with 100 bootstrap replications^30^. Finally, to gain additional confidence in the biochemical function of the proteorhodopsin genes, we analysed the primary amino acid sequence for the presence of proton pumping and light absorbing motifs^21^.

### *P. arcticus* 16S rRNA time-series

Samples were collected during the Joint Ocean Ice Study cruise to the Canada Basin between 2017-2023. For each sample, 4–14 L of seawater was sequentially filtered through a 3 μm pore size polycarbonate filter and a 0.22 μm pore size Sterivex filter (Durapore; Millipore, Billerica, MA, USA). Filters were preserved in RNAlater and stored at −80 °C until processed in the laboratory. DNA was extracted from the Sterivex filter using the following method: filters were thawed on ice and RNAlater storage buffer was removed and added to a 4 mL Amicon Ultracentrifugal filter unit (30 kDa). The Sterivex filter was washed with 1.5 mL of a sucrose-based lysis buffer rotated briefly and added to the Amicon. The Amicon was centrifuged at 7500 x g for 10-20 min. Buffer exchange was performed twice in the Amicon by addition of 2 mL lysis buffer. The storage buffer concentrate was added back to the Sterivex along with an additional volume of the lysis buffer to adjusting the total volume in the Sterivex to 1.4 mL. Filters were treated with 100 μL of 125 mg mL^−1^ lysozyme and incubated with rotation at 37 °C for 30 min. After incubation, 100 μL of 10 mg mL^−1^ proteinase K and 100 μL of 20% SDS was added. Filters were left to rotate for 2 h at 55 °C. Following cell lysis and digestion the lysate was collected from the filters and treated with 0.583 volumes of MPC Protein Precipitation Reagent (Epicentre, Madison, WI, USA) for and centrifugation at 10,000×g at 4 °C for 10 min. The supernatant was transferred to a clean tube. DNA was precipitated overnight at –20 °C with 1:10 volume of 3M sodium acetate and 2.5 volumes of cold 99-100% ethanol. Following centrifugation at 16,000 x g at 4 °C for 30 minutes, supernatant was removed, washed with cold 70% ethanol and resuspended in TE (pH8).

We used the 515-Y forward primer (5’-GTG YCA GCM GCC GCG GTA A –3’) from^38^, and the 806-RB reverse primer (5’-GGA CTA CNV GGG TWT CTA AT –3’) from^39^. PCR reactions (20 μL) contained 0.5 μM of each primer and 10 μL Phusion Hot Start II High-Fidelity PCR Master Mix (Thermo Scientific). Cycling conditions were as follows i) an initial 30 second denaturing step at 98°C, followed by ii) 30 cycles consisting of a 10 second annealing step at 98°C and a 30 second step at 50°C, and a 30 second extension of 30 seconds at 72°C, iii) and a final elongation step of 5 minutes at 72°C. The PCR amplicons, with the i5 and i7 adapters attached, were purified at Centre CERMO-FC (UQÀM) and subsequently sequenced using the Illumina MiSeq 250bp paired-end platform. Raw reads were processed using the DADA2 package in R^40^. Quality profiles were created for all reads. Reads were then trimmed via *cutadapt* in RStudio, merged and dereplicated, followed by ASV identification and chimera removal. We used blastn^41^ to identify the amplicon sequence variant (ASV) which represented *P. arcticus* within the Arctic Ocean 16S rRNA ASV time series.

Sea ice concentrations for the coordinates of each station (BL8, CB21, CB4, CB9, ICE) resolved within 25 km^2^, were obtained from the National Snow and Ice Data Center database. Daily sea ice concentration data was retrieved for each sampling location, from the date of sampling up to 4 months before collection, via RStudio access of the NSIDC dataset G02135.

## Data availability

Single sample metagenome and metatranscriptome dataset are available at the Joint Genome Institute Genomes OnLine Database under study ID Gs0134626 (see **Table S1** for details). Two co-assemblies are also available under study ID Gs0134626: the CB co-assembly (Analysis ID Ga0485091) and the Upper water co-assembly (Analysis ID Ga0485101). The western Arctic Ocean MAG catalog is available at Dryad (https://doi.org/10.5061/dryad.g4f4qrfxk). The *P. arcticus* genome is available at NCBI (JAZHGD000000000; BioSample SAMN39732788). Metagenome sequence data is available through NCBI (PRJNA539549,PRJNA539628,PRJNA539629,PRJNA539670,PRJNA539671,PRJNA539672,PRJNA539673,PRJNA539674,PRJNA539675,PRJNA539676,PRJNA539677,PRJNA539678,PRJNA539679,PRJNA539680,PRJNA539681,PRJNA539682,PRJNA539683,PRJNA539684,PRJNA539685,PRJNA539686,PRJNA539687,PRJNA539688,PRJNA539689,PRJNA539690,PRJNA539691,PRJNA539692,PRJNA539693,PRJNA539694,PRJNA622038,PRJNA622039,PRJNA 622040) Metatranscriptome sequence data is available through NCBI (PRJNA520201,PRJNA520202,PRJNA520206,PRJNA520208,PRJNA520210,PRJNA520212,PRJNA520215,PRJNA520218,PRJNA520220,PRJNA520222,PRJNA520224,PRJNA520228,PRJNA520231,PRJNA520233,PRJNA520235,PRJNA520238,PRJNA520240,PRJNA520243,PRJNA520254,PRJNA520257,PRJNA520260,PRJNA520262,PRJNA535976,PRJNA535977,PRJNA535978,PRJNA535979,PRJNA535980,PRJNA537841,PRJNA537842,PRJNA537843) Source data are provided with this paper.

## Acknowledgments

The data were collected aboard the CCGS Louis S. St-Laurent in collaboration with researchers from Fisheries and Oceans Canada at the Institute of Ocean Sciences and Woods Hole Oceanographic Institution’s Beaufort Gyre Exploration Program and are available at http://www.whoi.edu/beaufortgyre. We would like to thank both the captain and crew of the CCGS Louis S. St-Laurent and the scientific teams aboard. The work was conducted in collaboration with “Facilities Integrating Collaborations for User Science” (FICUS) program between the Joint Genome Institute (JGI) and the Environmental Molecular Sciences Laboratory (EMSL).

## Authors Contributions Statement

S.L. and D.A.W. conceived the study, designed the work, and interpreted the data. S.L., S.P., and A.W. performed the bioinformatic analyses. T.G. performed the *P. hydrocarbonoclasticus* growth experiments. S.L. and D.A.W wrote the manuscript. All authors read and revised the manuscript.

