## Supplementary Material for "Evidence of an acetone carboxylation pathway in photoheterotrophic bacteria from the Arctic Ocean"

**McLatchie et al. Supplementary Materials**

### Supplementary Methods

#### Metagenomic assembly and co-assembly

Details of seawater sampling, DNA extraction, shotgun sequencing, and generation of single sample metagenome assemblies were previously described in[1]. In the current study, we generated metagenome co-assemblies of collection of samples from the Canada Basin and Amundsen Gulf (**Supplementary Data 1**) to recover a greater diversity of MAGs. All metagenome co-assemblies were generated using MEGAHIT v.1.2.7[2] with k-mer sizes of 27, 37, 47, 57, 67, 77, and 87.

#### Binning, dereplication, and taxonomic classification of MAGs

The input metagenome reads for each co-assembly were mapped to the co-assemblies with BWA v.0.7.17 using the mem option[3]. The mapping results were processed using jgi_summarize_bam_contig_depths. Contig binning was performed for each of the assemblies using MetaBAT2 v.2.12.1[4] with default settings, resulting in 4,824 genome bins. Genome quality was evaluated using CheckM v.1.0.11[5] with the lineage_wf workflow. The 924 bins with >50% completeness and <10% contamination and strain heterogeneity were considered at least medium-quality MAGs. Dereplicated MAGs with >50% completeness and <10% contamination and strain heterogeneity were taxonomically classified with the GTDB-tk v.1.3.0 using the classify_wf workflow[6]. The medium-quality MAGs were dereplicated with dRep using 95% ANI cut-off to remove species level redundancy[7], resulting in 663 representative MAGs.

#### MAG abundances and distributions across the global ocean

To determine the coverage of MAGs in the Canada Basin metagenomes (**Supplementary Data 1**) and 129 TARA global oceans metagenomes[8] and 20 Southern Ocean metagenomes[9], trimmed metagenome reads were initially recruited to a concatenation of the MAGs at a minimum sequence identity of 98% using BBMap v.35. The mapping files were parsed to extract the mapping information for each individual MAG using a custom Unix script which implemented SAMtools[10]. Mapping files for each MAG were converted from SAM to sorted BAM format files using SAMtools[10]. Horizontal coverage was calculated following the method in[11] in which the average sequencing depth is truncated to the central 80% of the mapped positions ($TAD_{80}$) normalized by the number of genome equivalents. The number of genome equivalents in each metagenome was estimated using MicrobeCensus v.1.1.0[12]. A principal coordinate analysis was performed to assess the community composition of medium-quality MAG assemblages based on the Bray-Curtis dissimilarities in $TAD_{80}$ abundances using the function pcoa() in the R package vegan[13].

#### Functional annotation of MAGs

Gene prediction and annotation was performed using Prokka v.1.12[14], which implemented Prodigal v.2.6.3 for protein coding genes[15], Barrnap for ribosomal RNA genes, and ARAGORN v.1.2[16] for transfer and transfer-messenger RNA genes. Functional annotation of proteins was performed using Prokka v.1.12[14] with default settings. Functional annotation of proteins was also performed using KofamKOALA[17] with default settings and a threshold score greater than 0.7 or higher and an e-value below $1 \times 10^{10}$ or lower to identify metabolic genes within the MAG catalogue.

### Supplementary Results

#### The Western Arctic Ocean MAG catalogue

We generated a MAG catalogue from an eight station transect of the western Arctic Ocean that included the Amundsen Gulf and a latitudinal gradient of the Canada Basin (73-81°N) (**Figure S1a**). We performed a metagenomic survey of the oligotrophic Canada Basin samples (Table S1). All stations were ice-free during the sampling period (September 2017), except for the most northern station (CBN3). The survey targeted distinct water layers, including the surface mixed layer (SML; 5 m and 20 m), the subsurface chlorophyll maximum (SCM; 55-95 m), Pacific-origin winter water (PWW; 90-250 m) and Atlantic-origin water (AW; 360-1000 m) (**Figure S1b**). To maximize the MAG diversity recovered, we performed contig binning on each of the 31 single metagenome assemblies, as well as on three metagenome coassemblies generated from 1) all 24 Canada Basin samples (CB coassembly), 2) 11 Canada Basin samples from the SML and SCM (Upper water coassembly), and 3) 7 Amundsen Gulf samples (AG coassembly). In total, we recovered 924 MAGs exceeding MIMAG medium quality through an additional <10% strain heterogeneity criterion. MAGs were dereplicated across assemblies/ coassemblies (average nucleotide identity >= 95%), resulting in a genomospecies-level set of 663 MAGs. Of the genomospecies, 84% (559) were represented by MAG originating from metagenome coassemblies and 16% (104) originated from single metagenome assemblies (**Supplementary Data 2**), demonstrating the value of combining coassembly and single assemblies in generation of MAG datasets.

To map MAG biogeography across the water layers of the Canada Basin, we calculated the central 80% truncated average depths of coverage ($TAD_{80}$) from fragment recruitment. Based on $TAD_{80}$, between 62 and 471 MAGs were detected across samples, with a pattern of increased observed richness with depth (**Figure S1c**). Correspondingly, fragment recruitment to the MAG dataset was lowest in the surface waters (average 8.34%) and increased with depth with the highest mean read recruitment from the PWW samples (26.34%), indicating less genomic representation from the surface waters (**Figure S1d**) A principal coordinate analysis demonstrated distinct MAG assemblages were strongly structured by the stratified waters masses of the Western Arctic Ocean (**Figure S1e**).

We next investigated the distribution of the Arctic Ocean MAG catalogue across other regions of the global ocean. The Arctic Ocean MAG catalogue recruited on average 0.05% (0.04% and 0.08%) of reads from surface and chlorophyl maximum layer metagenomes from other regions of the ocean (**Figure S1d)**. Recruitment was greater for mesopelagic metagenomes from lower latitudes (0.14 to 6.19 %) and was greatest for a single mesopelagic metagenome from the Southern Ocean (9.74 %). This restricted distribution across global oceans demonstrate a strong Arctic Ocean endemism of the MAGs.

The MAG dataset represented a wide phylogenetic range, including 619 bacterial (**Figure S2a**) and 44 archaeal taxa (**Figure S2b**) of which the majority (93%) were novel at the species level (**Figure S2c**).

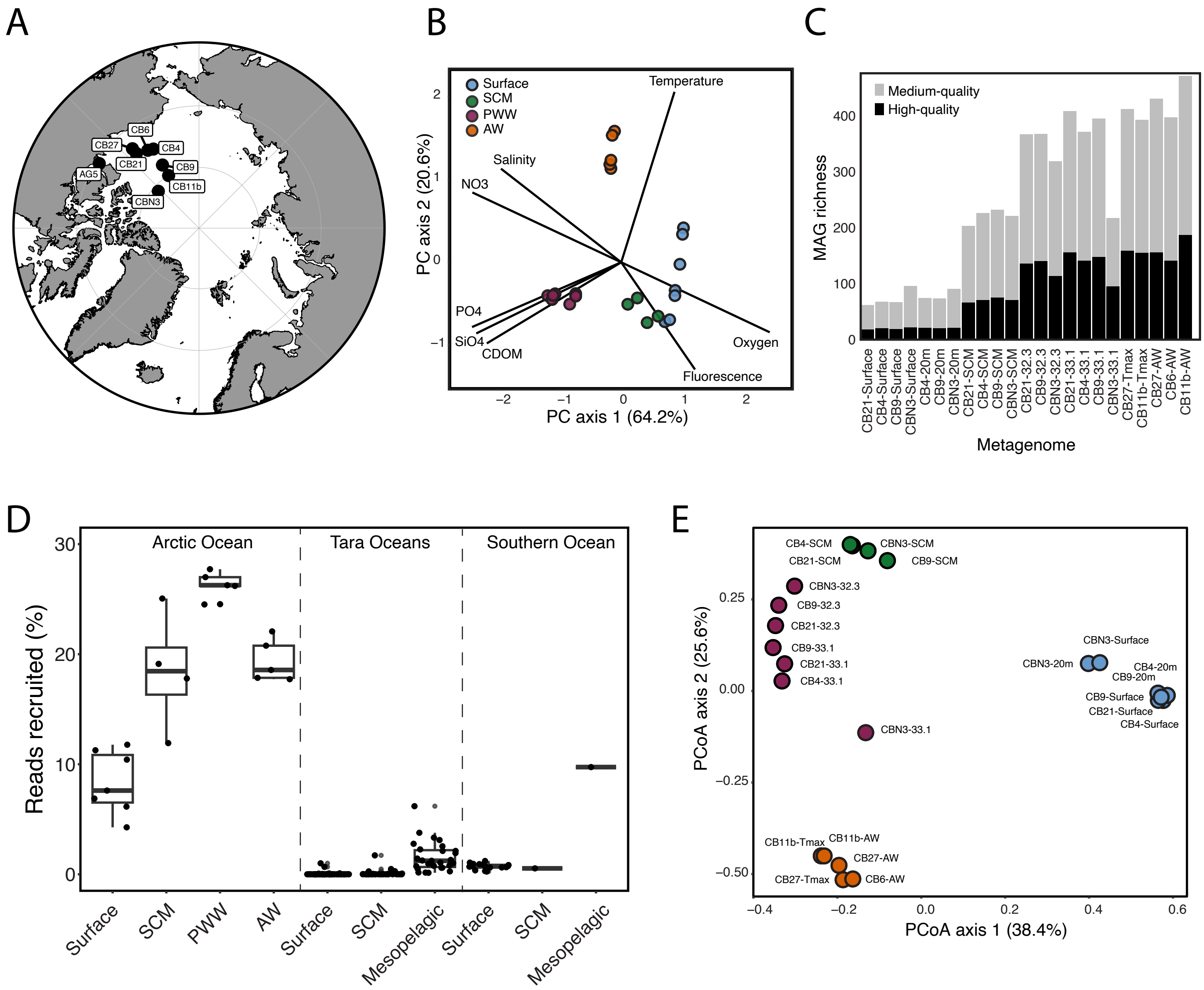
A
CB6
CB27
CB4
CB21
CB9
AG5
CB11b
CBN3
B
Surface
SCM
PWW
AW
Temperature
Salinity
NO3
PO4
SiO4
CDOM
Oxygen
Fluorescence
PC axis 1 (64.2%)
PC axis 2 (20.6%)
C
Medium-quality
High-quality
MAG richness
Metagenome
CB21-Surface
CB4-Surface
CB9-Surface
CBN3-Surface
CB4-20m
CB9-20m
CBN3-20m
CB21-SCM
CB4-SCM
CB9-SCM
CBN3-SCM
CB21-32.3
CB9-32.3
CBN3-32.3
CB21-33.1
CB4-33.1
CB9-33.1
CBN3-33.1
CB27-Tmax
CB11b-Tmax
CB27-AW
CB6-AW
CB11b-AW
D
Arctic Ocean
Tara Oceans
Southern Ocean
Reads recruited (%)
Surface
SCM
PWW
AW
Surface
SCM
Mesopelagic
Surface
SCM
Mesopelagic
E
CB4-SCM
CBN3-SCM
CB21-SCM
CB9-SCM
CBN3-32.3
CB9-32.3
CB21-32.3
CB9-33.1
CB21-33.1
CB4-33.1
CBN3-33.1
CBN3-Surface
CBN3-20m
CB4-20m
CB9-20m
CB9-Surface
CB21-Surface
CB4-Surface
CB11b-Tmax
CB11b-AW
CB27-AW
CB27-Tmax
CB6-AW
PCoA axis 1 (38.4%)
PCoA axis 2 (25.6%)

**Figure S1.** Vertical structuring of the microbial community in the Canada Basin, Arctic Ocean. **(A)** Map of sampling stations in the Canada Basin, Arctic Ocean. The map was generated using ggOceanMaps in R. **(B)** Principal component analysis of the environmental characteristics of the Canada Basin metagenome sampling locations. **(C)** The number of MAGs recovered form metagenome assemblies. **(D)** Abundance of the MAG collection in the Arctic Ocean and global oceans based on fragment recruitment **(E)** Principal coordinate analysis of the taxonomic variation among Arctic Ocean MAG assemblages based on $TAD_{80}$ community fraction of the MAGs.

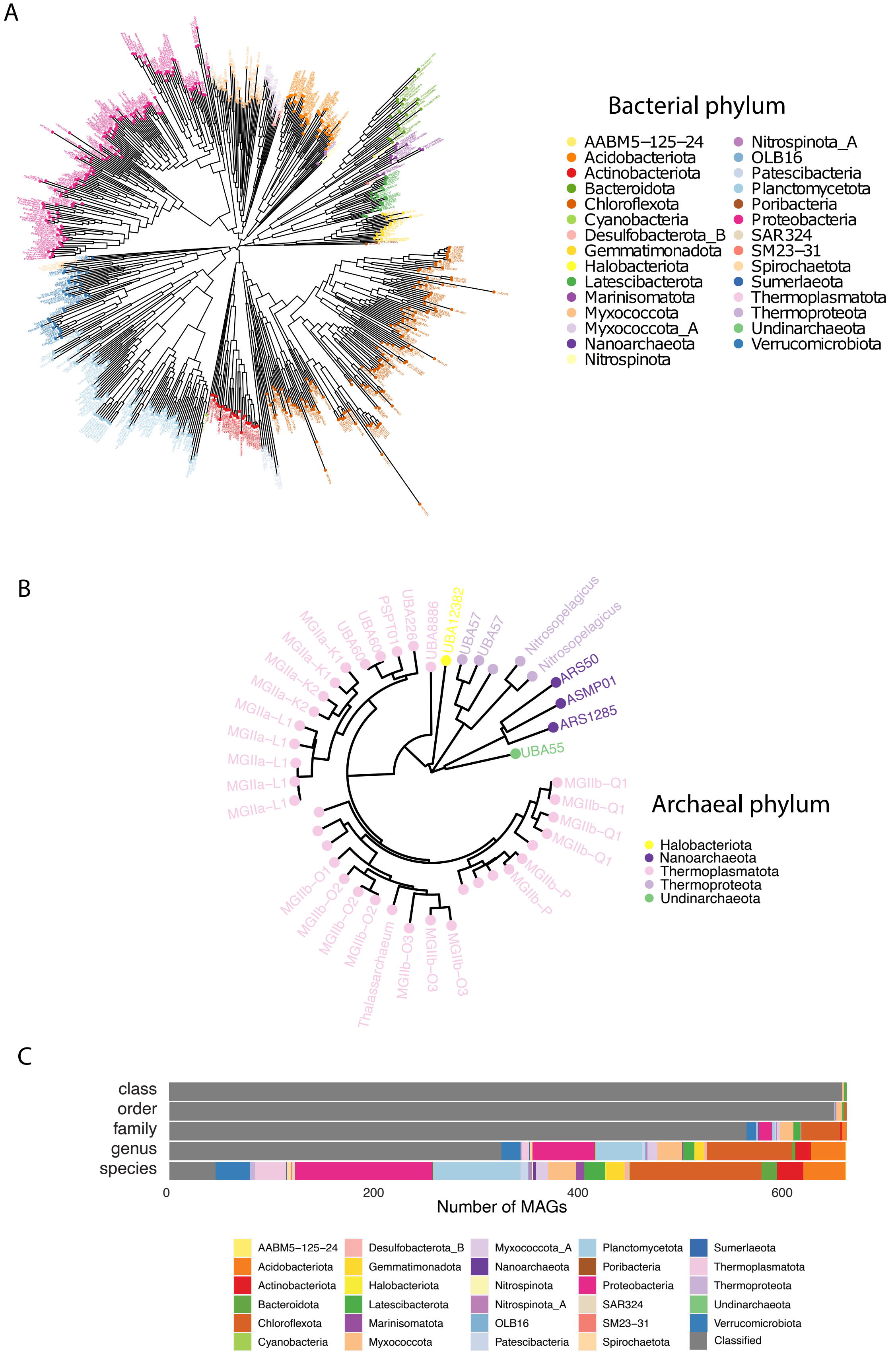
A
Bacterial phylum
AABM5−125−24
Acidobacteriota
Actinobacteriota
Bacteroidota
Chloroflexota
Cyanobacteria
Desulfobacterota_B
Gemmatimonadota
Halobacteriota
Latescibacterota
Marinisomatota
Myxococcota
Myxococcota_A
Nanoarchaeota
Nitrospinota
Nitrospinota_A
OLB16
Patescibacteria
Planctomycetota
Poribacteria
Proteobacteria
SAR324
SM23−31
Spirochaetota
Sumerlaeota
Thermoplasmatota
Thermoproteota
Undinarchaeota
Verrucomicrobiota
B
MGIIa−K1
MGIIa−K1
MGIIa−K2
MGIIa−K2
MGIIa−L1
MGIIa−L1
MGIIa−L1
MGIIa−L1
MGIIa−L1
UBA60
UBA60
PSPT01
UBA226
UBA8886
UBA12382
UBA57
UBA57
Nitrosopelagicus
Nitrosopelagicus
ARS50
ASMP01
ARS1285
UBA55
MGIIb−Q1
MGIIb−Q1
MGIIb−Q1
MGIIb−Q1
MGIIb−P
MGIIb−P
MGIIb−O1
MGIIb−O2
MGIIb−O2
MGIIb−O2
Thalassarchaeum
MGIIb−O3
MGIIb−O3
MGIIb−O3
Archaeal phylum
Halobacteriota
Nanoarchaeota
Thermoplasmatota
Thermoproteota
Undinarchaeota
C
class
order
family
genus
species
0
200
400
600
Number of MAGs
AABM5−125−24
Acidobacteriota
Actinobacteriota
Bacteroidota
Chloroflexota
Cyanobacteria
Desulfobacterota_B
Gemmatimonadota
Halobacteriota
Latescibacterota
Marinisomatota
Myxococcota
Myxococcota_A
Nanoarchaeota
Nitrospinota
Nitrospinota_A
OLB16
Patescibacteria
Planctomycetota
Poribacteria
Proteobacteria
SAR324
SM23−31
Spirochaetota
Sumerlaeota
Thermoplasmatota
Thermoproteota
Undinarchaeota
Verrucomicrobiota
Classified

**Figure S2. A MAG catalogue from the western Arctic Ocean.** Phylogenetic diversity of (**A**) bacterial and (**B**) archaeal MAGs in the Western Arctic Ocean MAG catalogue. For the archaeal tree, taxa IDs were not assigned for some MAGs at this taxonomic level. (**C**) Phylogenetic novelty of the MAGs. The bars represent the number of MAGs within that taxonomic group that are novel at each of the five taxonomic levels.

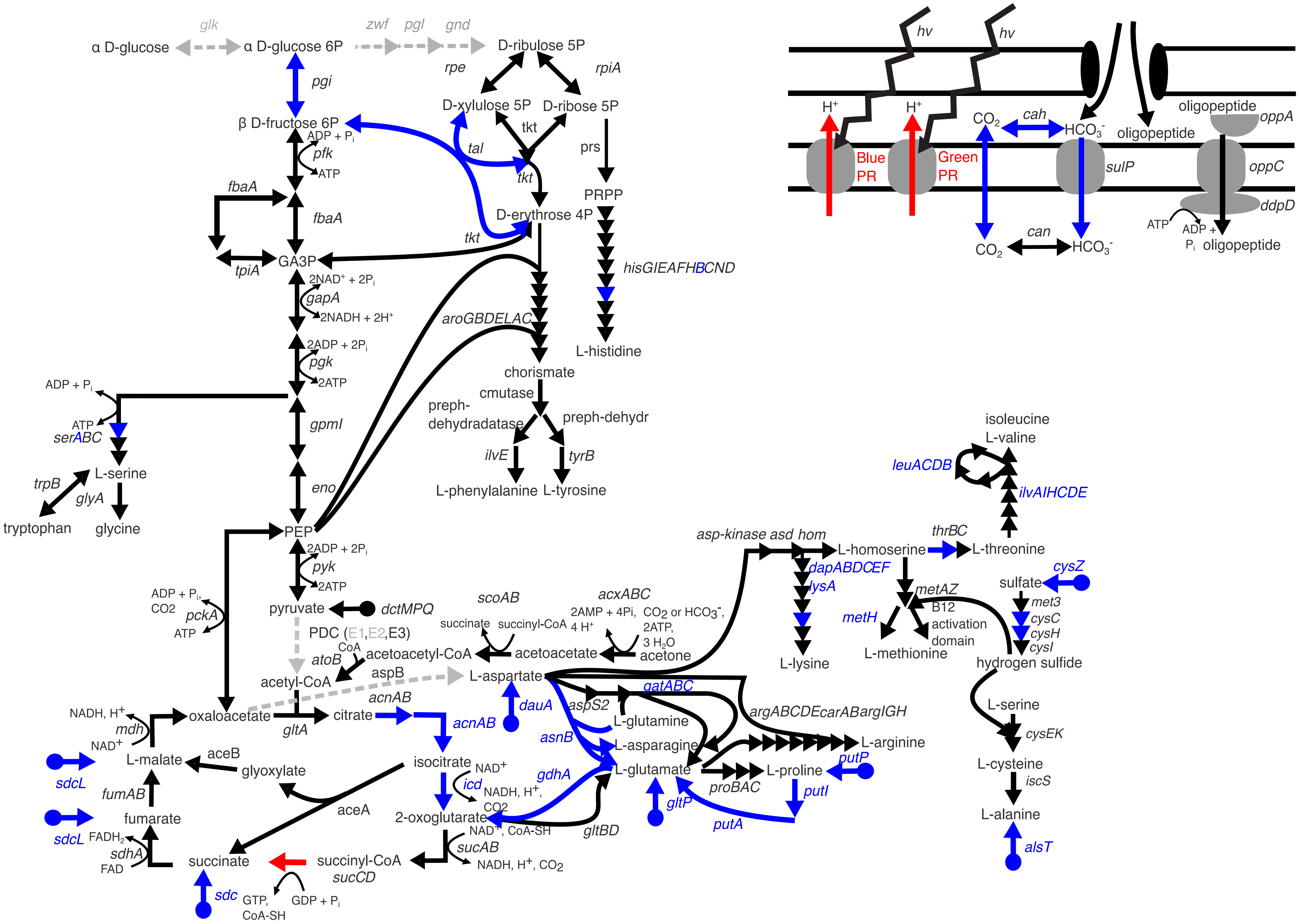

α D-glucose
glk
α D-glucose 6P
zwf
pgl
gnd
D-ribulose 5P
rpe
rpiA
pgi
D-xylulose 5P
D-ribose 5P
β D-fructose 6P
ADP + Pi
pfk
ATP
tal
tkt
prs
PRPP
fbaA
D-erythrose 4P
tpiA
GA3P
2NAD+ + 2Pi
gapA
2NADH + 2H+
hisGIEAFHBCND
aroGBDELAC
L-histidine
2ADP + 2Pi
pgk
2ATP
chorismate
cmutase
preph-dehydradatase
preph-dehydr
gpmI
serABC
ilvE
tyrB
L-serine
trpB
glyA
L-phenylalanine
L-tyrosine
tryptophan
glycine
eno
PEP
pyk
pyruvate
dctMPQ
ADP + Pi,
CO2
pckA
ATP
PDC (E1,E2,E3)
scoAB
acxABC
succinate
succinyl-CoA
2AMP + 4Pi,
4 H+
CO2 or HCO3-,
2ATP,
3 H2O
acetone
CoA
atoB
acetoacetyl-CoA
acetoacetate
acetyl-CoA
aspB
L-aspartate
asp-kinase asd hom
L-homoserine
thrBC
L-threonine
dapABDCEF
lysA
L-lysine
metAZ
metH
B12 activation domain
L-methionine
isoleucine
L-valine
leuACDB
ilvAIHCDE
cysZ
sulfate
met3
cysC
cysH
cysI
hydrogen sulfide
L-serine
cysEK
L-cysteine
iscS
L-alanine
alsT
NADH, H+
mdh
NAD+
oxaloacetate
gltA
citrate
acnAB
dauA
aspS2
gatABC
L-glutamine
asnB
L-asparagine
argABCDEcarABargIGH
L-arginine
putP
L-glutamate
gdhA
proBAC
L-proline
putI
gltP
putA
sdcL
L-malate
aceB
glyoxylate
isocitrate
NAD+
icd
NADH, H+,
CO2
fumAB
aceA
2-oxoglutarate
gltBD
fumarate
FADH2
sdhA
FAD
NAD+, CoA-SH
sucAB
NADH, H+, CO2
succinate
succinyl-CoA
sucCD
sdc
GTP,
CoA-SH
GDP + Pi
hv
hv
H+
H+
Blue PR
Green PR
CO2
cah
HCO3-
oligopeptide
oppA
oligopeptide
sulP
oppC
ddpD
can
CO2
HCO3-
ATP
ADP + Pi
oligopeptide

**Figure S3.** Metabolic reconstruction of the central carbon metabolism of *P. arcticus* and *P. hydrocarbonoclasticus*. Black arrows indicate genes of pathways conserved between *P. arcticus* and *P. hydrocarbonoclasticus*. Red arrows indicate genes of pathways possessed only by *P. arcticus*. Blue arrows indicate genes of pathways possessed only by *P. hydrocarbonoclasticus*. Grey dashed arrows indicate genes of pathways that are absent in *P. arcticus* and *P. hydrocarbonoclasticus*.
